# Post-transcriptional Regulation is the Major Driver of microRNA Expression Variation

**DOI:** 10.1101/2020.01.07.897975

**Authors:** Dingyao Zhang, Jingru Tian, Christine Roden, Jun Lu

**Author notes:** Correspondence should be addressed to: Jun Lu, 10 Amistad Street, Room 237C, New Haven, CT 06520.

## Abstract

MicroRNA (miRNA) expression patterns are highly variable across human tissues and across cancer specimens. The intuitive assumption is that transcription is the main contributor to mature miRNA expression patterns, with post-transcriptional processes further modifying miRNA expression levels. Here we report the surprising model that, on the global level, post-transcriptional regulation dominates over transcriptional regulation in determining mature miRNA expression patterns in both normal tissues and cancer. Taking advantage of large genomic datasets in which the expression of both mature miRNAs and their host genes have been quantified, we establish and validate transcriptional and post-transcriptional metrics, with miRNA host gene expression estimating transcriptional regulation and mature miRNA to host gene ratio estimating post-transcriptional regulation. On average, the post-transcriptional metric contributes 2.8-fold more than the transcriptional metric to the variance of mature miRNA expression. The variation of the balance between the two mature miRNAs (5p and 3p miRNAs) produced from the same precursor hairpin is a non-negligible contributor to miRNA expression, explaining ∼27% of the variance of miRNAs’ post-transcriptional metric. Data of normal tissues yield similar results as cancer specimens. Additionally, the post-transcriptional metric is superior to the transcriptional metric in classifying cancer types. We further demonstrate that the post-transcriptional metric separates miRNAs into distinct groups, suggesting that there are groups of miRNAs that are co-regulated on the post-transcriptional level. Our data support a model in which the post-transcriptional regulation is the major driver of miRNA expression variation, and paves a way toward better mechanistic understanding of post-transcriptional regulation of mature miRNA expression.

## Introduction

MicroRNAs (miRNAs) are small (∼22-nucleotide) non-coding RNAs that post-transcriptionally control cellular gene expression through translational inhibition and/or mRNA degradation ^1, 2, 3^. In normal and cancer specimens, miRNA expression levels vary substantially both across tissue types and across individual cancer specimens in the same cancer ^4, 5, 6, 7^. This property of miRNA expression variation could be utilized to classify human cancers, and in some cases, miRNA expression performs better than genome-wide mRNA expression in cancer classification ^4^. However, on the global level, how the variation of miRNA expression in cancer is regulated is not clear.

The first step of miRNA biogenesis involves the transcription of long hairpin-containing primary miRNA transcript (pri-miRNAs) by RNA polymerase II ^3, 8, 9^. Since human cancers display large variations in epigenetic states and often harbor mutations in epigenetic enzymes ^10^, it is reasonable to assume that transcriptional regulation drives the majority of miRNA expression variation in cancer. Following transcription, the vast majority of human miRNAs follow the canonical miRNA biogenesis pathway ^11^, in which DROSHA processes pri-miRNAs into precursor miRNAs (pre-miRNAs), which is further processed by DICER to generate miRNA duplexes involving two miRNA sequences (5p and 3p miRNA respectively) from each pre-miRNA hairpin ^12^. Loading of the 5p and 3p miRNAs into Argonaute (AGO) proteins results in functional mature miRNAs in cells which can further undergo regulations at the levels of stability and recycling ^13, 14, 15, 16^. Deregulation of mature miRNA expression at the post-transcriptional stages has been reported, which is predominantly elucidated at the steps of DROSHA- and DICER-mediated processing ^9, 12, 17^. However, how much miRNA expression variation in cancer is controlled transcriptionally and how much on the post-transcriptional level are not clear. We set out to address this question and to determine how miRNA expression variation is regulated.

## Results

To estimate the transcriptional versus post-transcriptional regulation of miRNA expression in cancer, we established a bioinformatics procedure to utilize public genomic datasets (such as The Cancer Genome Atlas (TCGA) ^18^ and Therapeutically Applicable Research To Generate Effective Treatments (TARGET) ^19^) datasets in which both small RNA and mRNA expression have been quantified across multiple cancer types. We took advantage of the fact that miRNAs often reside within defined host genes in the genome and are co-transcribed with their host genes. These include both miRNAs that are located within introns of the host genes and miRNAs that reside in exons of the host genes. Motivated by prior efforts exploiting relative abundance between expression levels of nascent messenger RNA (mRNA) and mature mRNA to estimate the rates of mRNA maturation ^20^, we reasoned that mature miRNA expression could be expressed as a multiplication product of transcriptional regulation and post-transcriptional regulation. Thus, we could use the miRNA host gene expression as a metric to estimate transcriptional regulation, and use the ratio between mature miRNA expression and the expression of the corresponding host gene (miRNA/host ratio) as a metric to estimate post-transcriptional regulation (Figure 1A). Indeed, our previous use of the miRNA/host ratio (i.e. post-transcriptional metric) shows that it faithfully reflects rules of pri-miRNA processing ^19, 21^.

**Figure 1.**
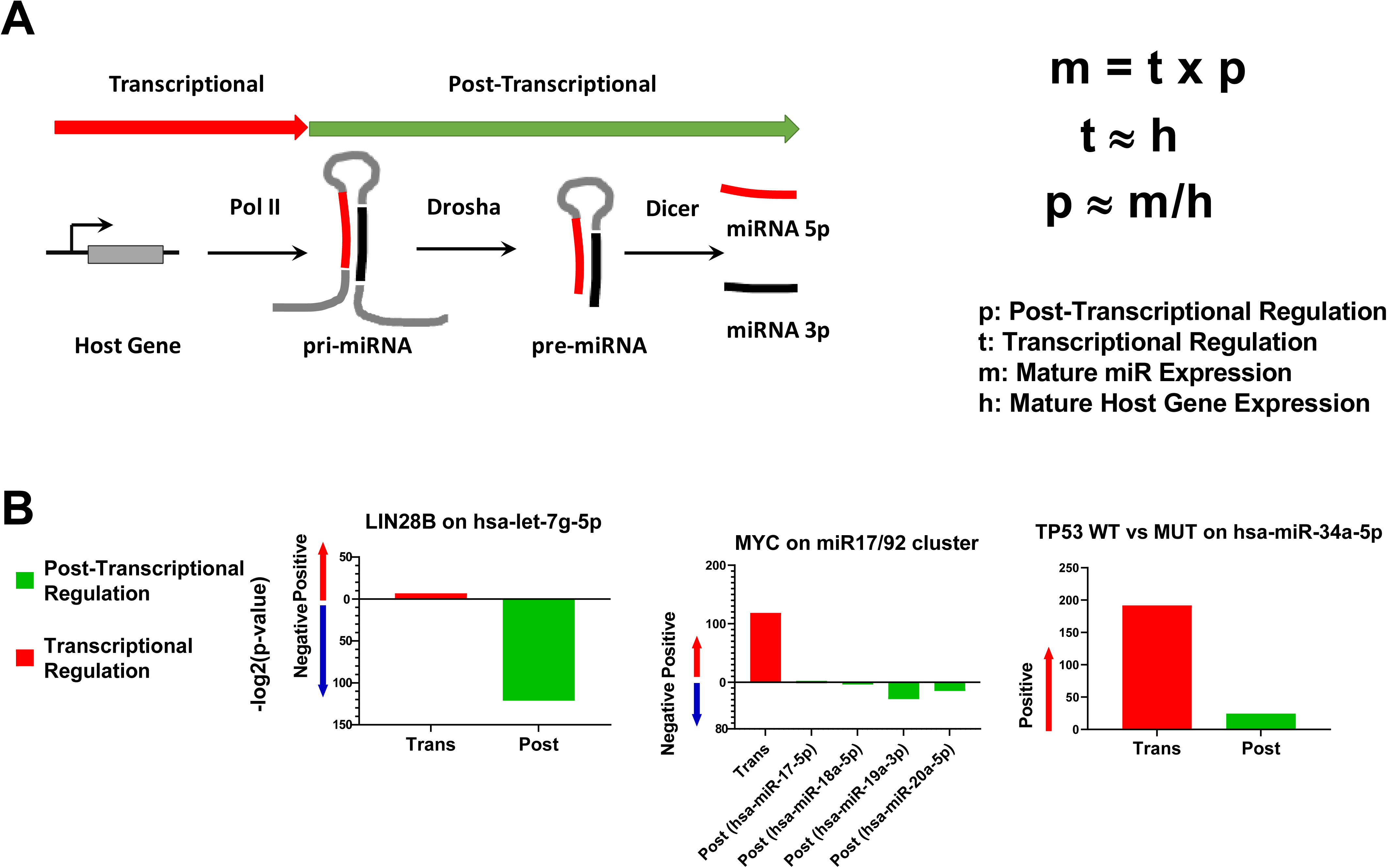
Development of transcriptional and post-transcriptional metrics on genomic datasets. **(A)** A schematic of canonical miRNA processing was shown on the left, with transcribed primary miRNAs (pri-miRNAs) processed sequentially by DROSHA and DICER into precursor miRNAs (pre-miRNAs) and mature miRNAs respectively. The two mature miRNAs from each miRNA hairpin are referred to as 5p and 3p miRNAs. Formulas used for estimating post-transcriptional regulation and transcriptional regulation are shown on the right. **(B)** Three cases for the verification of transcriptional and post-transcriptional metrics were plotted. Left: LIN28B-mediated suppression of let-7g processing, with significance of Spearman correlation on all TCGA cancer samples between LIN28B and the transcriptional and post-transcriptional metrics of hsa-let-7g-5p shown. Middle: c-Myc-mediated upregulation of miR-17/92 cluster transcription, similarly with significance of Spearman correlation shown; Right: p53 activation of miR-34a transcription, with TCGA samples separated into TP53 wildtype and mutant groups, and the significance of comparison between these two sample groups with regard to the transcriptional and post-transcriptional metrics of has-miR-34a-5p shown. Green bars reflect post-transcriptional metrics; red bars reflect transcriptional metrics.

We further determined that our transcriptional and post-transcriptional metrics could reflect known regulatory mechanisms of mature miRNA production. For example, for the known relationship of LIN28B-mediated suppression of the processing of *let-7g* ^22, 23, 24^, we observed in the TCGA dataset that *LIN28B* expression level is strongly anti-correlated with *let-7g*’s post-transcriptional metric (*let-7g*/host gene ratio) but not for *let-7g*’s transcriptional metric (*let-7g* host gene expression) (Figure 1B). For the known relationship of MYC regulating the transcription of miR-17/92 cluster ^25, 26^, we observed a strong positive correlation between *MYC* expression levels and the transcriptional metrics of miRNAs in the miR-17/92 cluster, but did not observe strong positive correlation with their post-transcriptional metrics (Figure 1B). Similarly, we examined the known relationship of p53 activating miR-34a transcription ^27, 28, 29, 30, 31^. Since it is well appreciated that p53 activity is heavily regulated on the protein level, we compared cancer specimens with p53 mutations to those that are wildtype, and observed that miR-34a’s transcriptional metric is strongly associated with *TP53* wildtype status, but much less so for miR-34a’s post-transcriptional metric (Figure 1B).

We next asked, on the global level, how much mature miRNA expression variation could be explained by transcriptional vs post-transcriptional metrics. Among the 2652 human mature miRNAs documented in miRBase release 22 ^32, 33^, we removed many that were expressed at undetectable or very low levels in nearly all samples in TCGA. To avoid ambiguity of host gene assignment, we further removed miRNAs that map to more than one locus or more than one host gene in the genome. After these filtering (see **Methods**), 813 mature miRNAs were retained for further analysis. For some of the miRNAs, such as hsa-miR-584-5p, we observed that the mature miRNA expression is strongly correlated with its transcriptional metric but not its post-transcriptional metric. For others, such as hsa-miR-140-5p, we observed a strong correlation with its post-transcriptional metric, but a much weaker correlation with its transcriptional metric (**Supplementary Figure S1**). We then evaluated how much of the variance of mature miRNA expression could be explained by transcriptional and post-transcriptional metrics, which could be determined by the coefficient of determination R square (R^2^) between mature miRNA expression and host gene expression or miRNA/host ratios. Surprisingly, when examining across 32 cancer types in TCGA, the transcriptional metric could only explain 18.2% of mature miRNA expression variation, whereas the post-transcriptional metric could explain 50.7% (∼2.8-fold more than the transcriptional metric, Figure 2A). When examining only within each individual cancer type, these numbers vary, but overall, the post-transcriptional metrics could explain the variance of mature miRNA expression >2-fold more than the transcriptional metrics (Figure 2A). Similar results could be observed using a different method of data normalization of the TCGA data (**Supplementary Figure S2**), and using an independent TARGET dataset (Figure 2A), thus arguing against the possibilities of dataset artifacts or normalization artifacts. Taken together, the data above support the notion that post-transcriptional regulation contributes more than transcriptional regulation to mature miRNA expression patterns.

**Figure 2.**
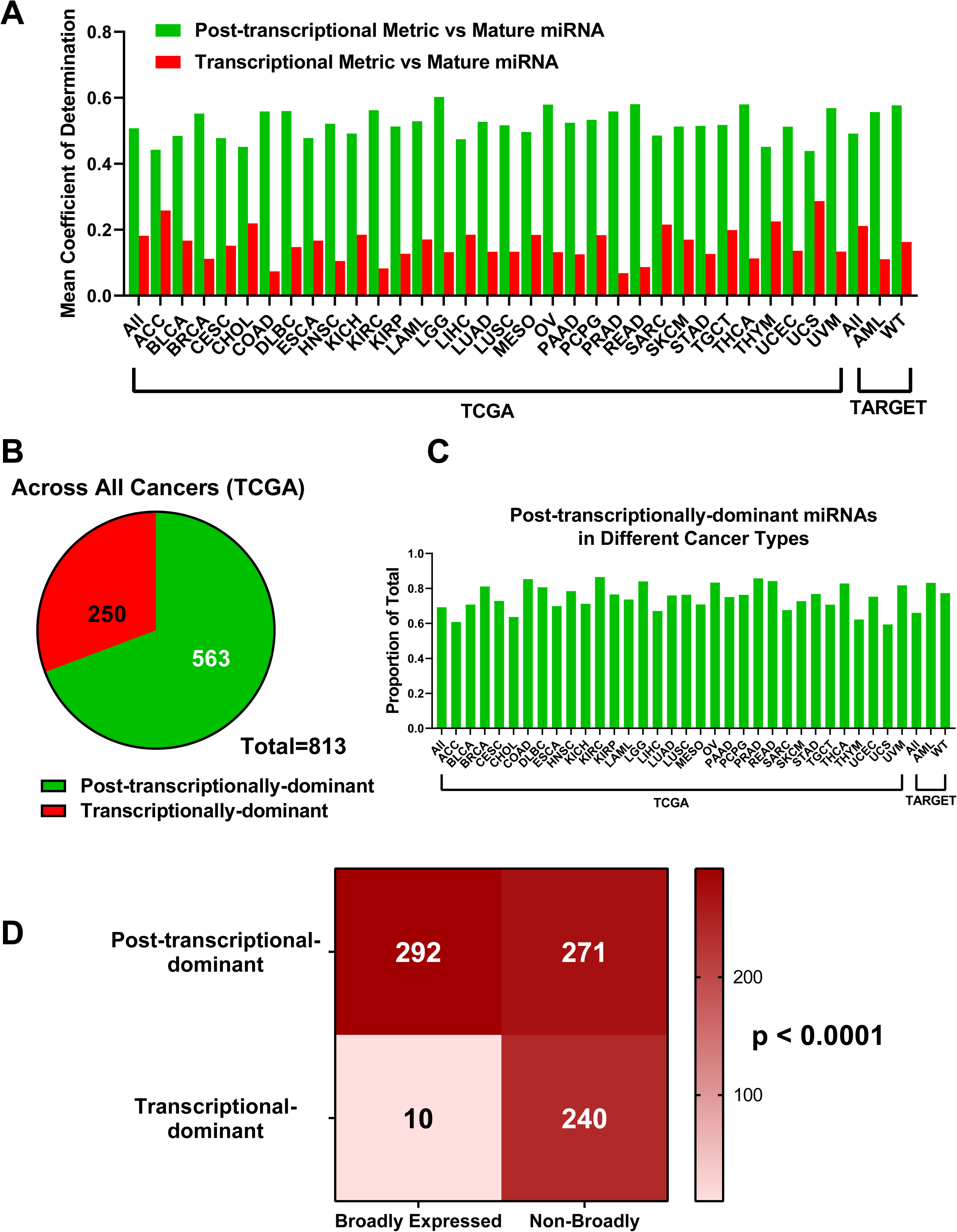
Post-transcriptional metrics contribute to the variation of mature miRNA expression more than transcriptional metrics in cancer samples. **(A)** A bar graph plotting average contribution of post-transcriptional metrics to overall miRNA expression variation (green bars) and average contribution of transcriptional metrics (red bars) in each cancer type or across all cancer samples is shown. Sample sources (TCGA and TARGET dataset) are indicated. **(B)** A pie chart showing the number of transcription-dominant miRNAs and post-transcription-dominant miRNAs, calculated across all cancer samples in the TCGA dataset. **(C)** Proportions of post-transcription-dominant miRNAs in indicated cancer types or across all cancer samples are shown. Sample sources (TCGA and TARGET dataset) are indicated. **(D)** miRNAs were separated into those with broadly-expressed host gene patterns and those with non-broadly-expressed host gene patterns. These two groups were further separated based on transcription-dominance or post-transcription-dominance. Numbers indicate the number of miRNAs, with colors reflecting the numbers. A color bar is shown on the right. P value based on Fisher exact test is indicated.

Specific miRNAs could differ in their major mode of regulations. To determine which miRNAs are predominantly regulated transcriptionally or post-transcriptionally, we calculated the coefficient of determination between each of mature miRNA and its transcriptional as well as post-transcriptional metrics. Across all TCGA samples, we observed that 250 (30.8%) of mature miRNAs revealed dominance by the transcriptional metric, whereas 563 (69.2%) of miRNAs showed dominance by the post-transcriptional metric (Figure 2B). Such a phenomenon is not a dataset-specific artifact, because we observed significant overlaps of post-transcription-dominant (and transcription-dominant) miRNAs among TCGA, TARGET and CCLE datasets (**Supplementary Figure S3**). When examining within individual cancer types in the TCGA dataset, we observed a similar phenomenon, with post-transcriptional metric being dominant for the majority (59.4%-86.5%) of miRNAs (Figure 2C). Additionally, the post-transcription-dominant miRNAs are not randomly distributed. We classified miRNA host genes based on their expression pattern into those that were broadly expressed across cancer specimens and those that had more restricted expression patterns (see **Methods**). We noticed that post-transcription-dominant miRNAs were preferentially enriched among those whose host genes showed broad expression patterns (see **Methods**) in comparison to those with more restricted host gene expression patterns (Figure 2D), suggesting the existence of a strategy to diversify mature miRNA expression for host genes that undergo less transcriptional regulation. These data are again consistent with post-transcriptional regulation being the main driver of miRNA expression variation.

While the data above support the notion that post-transcriptional regulation contributes more than transcription to shaping the landscape of miRNA expression variation, it remains possible that transcription control is more meaningful, whereas post-transcriptional regulation merely adds noise. To address this possibility, we first performed principle component analysis using the post-transcriptional metrics. We observed that specimens tend to cluster together according to cancer types (Figure 3A**, Supplemental Figure S4**). For example, acute myeloid leukemia (AML) specimens from both TCGA and TARGET are clustered in close proximity, despite these being two completely different datasets without undergoing batch correction (Figure 3A**, Supplemental Figure S4**). These data support that the post-transcriptional metric captures cancer-relevant information. To further confirm that the post-transcriptional metric encodes meaningful information with respect to cancer, we performed cancer type classification by using three categories of variables: the transcriptional metric, the post-transcriptional metric or mature miRNA expression level. We reasoned that if post-transcriptional regulation is dominantly informative, we would expect that the post-transcriptional metric would perform better in classification than the transcriptional metric. If not, we would expect the opposite (Figure 3B). Specifically, the TCGA dataset was randomly divided into training and test data, and support-vector machine (SVM)-based multi-class classifiers were developed on the training data for each variable category which were then used to predict the cancer types from the test dataset. Importantly, the number of variables in each variable category is the same to ensure fair comparison (**see Methods**). We observed that the classifier based on post-transcriptional metric performed significantly better than those from the transcriptional metric when evaluating Macro-F1 Scores (see **Methods**) (Figure 3C). We next asked whether post-transcriptional regulation could reflect cancer subtype information. We focused on breast cancer in TCGA, for which several histologic subtypes, including breast invasive ductal carcinoma, breast invasive lobular carcinoma as well as breast mixed ductal and lobular carcinoma, have been previously defined ^34, 35^. The post-transcriptional metric again outperformed the transcriptional metric in predicting these three histologic subtypes (Figure 3D). In contrast, breast cancer specimens could be grouped based on the similarity of DNA methylation patterns ^34, 35^, and we would expect that miRNAs’ transcriptional information is more effective at differentiating DNA methylation patterns which primarily impact transcription. Indeed, the transcriptional metric performed significantly better than the post-transcriptional metric when classifying the DNA methylation phenotypes (Figure 3E). Of note, performing predictions using a different filtering criterion yielded overall similar results (**Supplementary Figures S5A-S5C**). These results further validate the use of transcriptional and post-transcriptional metrics. Taken together, the data above support that miRNA post-transcriptional metric is superior to the transcriptional metric in classifying human cancer types, and further support the notion that post-transcriptional regulation is the major contributor to meaningful miRNA gene expression variation.

**Figure 3.**
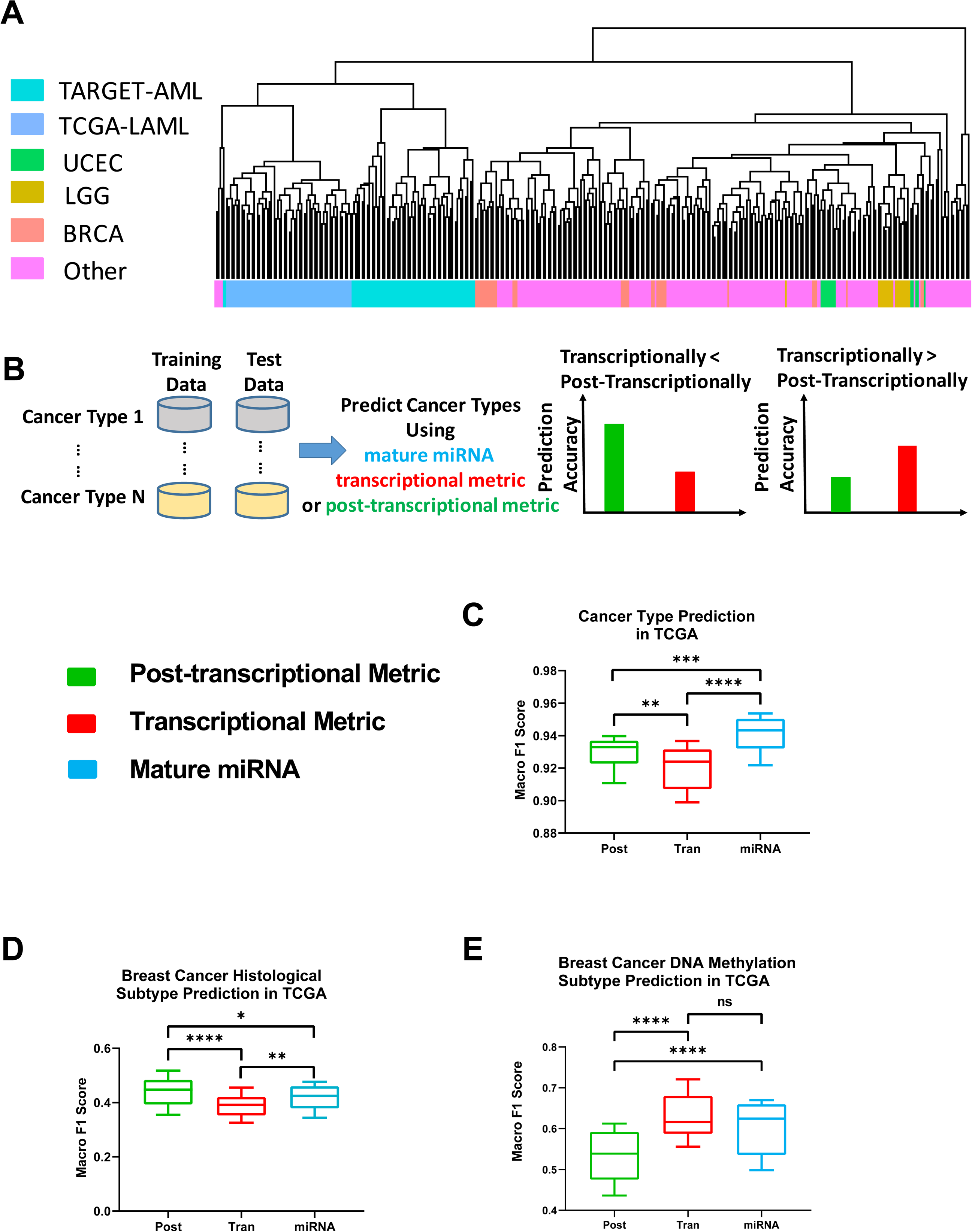
Post-transcriptional metrics are superior in cancer type prediction. **(A)** Hierarchical clustering was performed based on the post-transcriptional metrics of miRNAs in a randomly selected subset of TCGA samples (see Methods). The dendrogram from the clustering is shown with AML samples from TCGA and TARGET datasets indicated. Also indicated are the three largest TCGA cancer collections, including breast invasive carcinoma (BRCA), uterine corpus endometrial carcinoma (UCEC), and low grade glioma (LGG). A full version of the clustering result is available in Supplementary Figure S4. **(B)** A schematic showing the principles and pipelines of the prediction analyses. Transcriptional (red) and post-transcriptional (green) metrics were color coded, with blue showing predictions based on mature miRNA expression. **(C)** SVM-based multi-class predictors were developed based on the indicated metrics to predict cancer types in TCGA. Each test sample was classified into one of the 32 cancer types. miRNA/host gene pairs after filtering (see Methods) were used. Predication performance was plotted using box-and-whisker plot, with data representing Macro-F1 score results from multiple 10-fold cross-validation. **(D)** SVM-based multi-class predictors were developed to predict breast cancer histologic subtypes. Predication performance is shown. **(E)** SVM-based multi-class predictors were developed to predict breast cancer methylation subtypes. Predication performance is shown. (ns: not significant; * p <0.05; ** p < 0.01; *** p < 0.001; **** p < 0.0001).

To gain further insights into post-transcriptional processes that contribute to miRNA expression variation, we examined the balance between 5p and 3p miRNAs derived from the same hairpin, which is determined purely post-transcriptionally. In the canonical miRNA processing pathway, 5p and 3p miRNAs are produced at an 1:1 molar ratio though Dicer processing (Figure 1A) ^36^. Although the loading efficiency of 5p and 3p miRNAs into AGO proteins is often different, the current theory states that this efficiency difference is primarily determined by the sequence features of 5p and 3p miRNAs ^37, 38, 39^ (and thus there should not be major sample-based differences in AGO loading of 5p vs. 3p miRNAs), including the thermodynamic stability at the ends of the miRNA duplex ^38^ and the identity of the first nucleotide ^40^. An alternative pathway involves AGO2-mediated processing of pre-miRNAs, leading to selective incorporation of 5p miRNA into AGO2 ^41^. However, this alternative pathway is only required for the processing of one mammalian miRNA, i.e. miR-451 ^42, 43, 44^, and contributes <5% on average to the production of other mature miRNAs in human cell lines or zebrafish embryos ^11^. Based on these, one would not expect substantial levels of variation in miRNA 5p/3p ratios, and 5p and 3p miRNA expression should be highly correlated in cancer specimens. Surprisingly, when examining across all TCGA cancer datasets, we observed an average Spearman correlation coefficient of 0.516 between 5p and 3p miRNAs. For analysis within each cancer type, the average correlation coefficients ranged from 0.266 to 0.632, further supporting the existence of substantial variation in 5p/3p ratios. Similar results were observed in the TARGET dataset, arguing against the possibility of dataset-specific artifacts (Figure 4A). We also noticed that bimodal distributions of 5p/3p ratios existed for some miRNA, such as hsa-mir-146a which is an important tumor suppressor miRNA involved in hematopoietic stem cells and innate immune signaling ^45, 46^ (Figure 4B). To determine the contribution of miRNA 5p/3p ratio to overall miRNA post-transcriptional variation, we observed the coefficient of determination between 5p/3p miRNA ratios and the post-transcriptional metrics to be ∼0.27 across different cancer types (Figure 4C). Furthermore, miRNA 5p/3p ratios could classify human cancer types much better than random chance, albeit less accurately than post-transcriptional metrics (Figure 4D). These data argue for a non-negligible role of 5p-3p strand balance regulation in miRNA post-transcriptional regulation.

**Figure 4.**
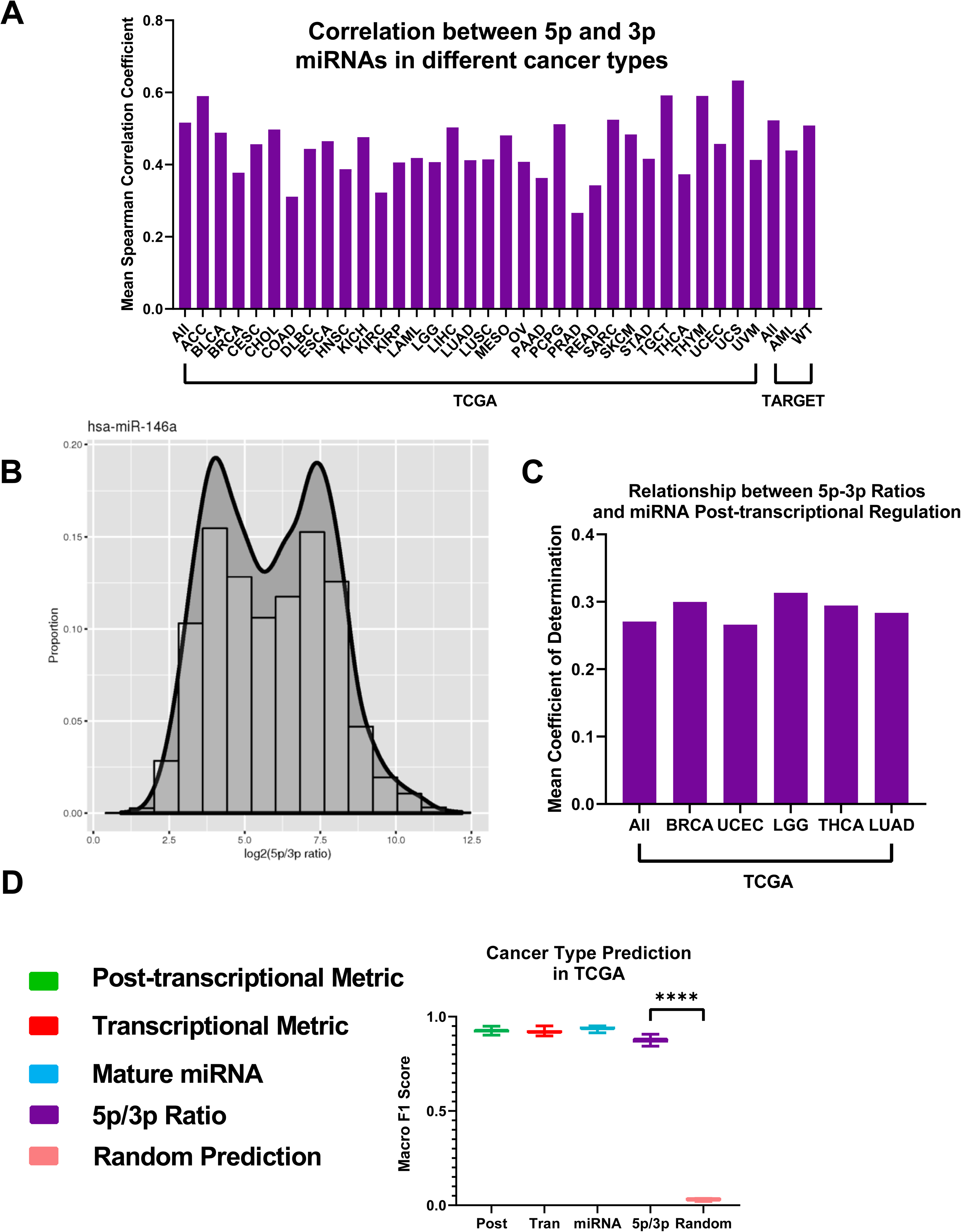
Variations of 5p and 3p miRNA ratios contribute to post-transcriptional variation. **(A)** A histogram showing mean Spearman correlation coefficients between 5p and 3p miRNAs (from the same hairpins) in each cancer type or across all cancer samples. Samples sources (TCGA or TARGET) are indicated. **(B)** A histogram of 5p/3p ratios of hsa-mir-146a was drawn along with a smoothed density estimate curve. **(C)** A histogram showing the contribution of the variance of the 5p/3p miRNA ratios to the variance of miRNA post-transcriptional metrics, across all cancer samples from TCGA dataset or within indicated cancer types. **D**. SVM-based multi-class predictors were developed based on the indicated metrics to predict cancer types in TCGA. Each test sample was classified into one of the 32 cancer types. Predication performance was plotted using box-and-whisker plot, with data representing Macro-F1 score results from multiple 10-fold cross-validation. (* p <= 0.05, ** p <= 0.01, *** p <= 0.001, **** p <= 0.0001)

Since the above analyses were performed on cancer samples, it is thus an interesting question whether similar phenomena occur in normal human tissues. We carried out similar analyses using normal specimens within TCGA. Similar to cancer specimens, the post-transcriptional metric contributed more to the variation of mature miRNA expression and showed stronger dominance in the variation of individual miRNAs than the transcriptional metric in normal specimens (Figure 5A, Figure 5B). The difference between post-transcriptional and transcriptional contributions seemed even more profound than that in cancer specimens, which might be partly due to the fact that only a few cancer types have sufficient normal specimens in TCGA. Also similar to cancer specimens, low correlation between 5p and 3p miRNAs was seen in normal samples (Figure 5C). These data support that the influence of post-transcriptional regulation on mature miRNA expression is not unique to the cancerous state.

**Figure 5.**
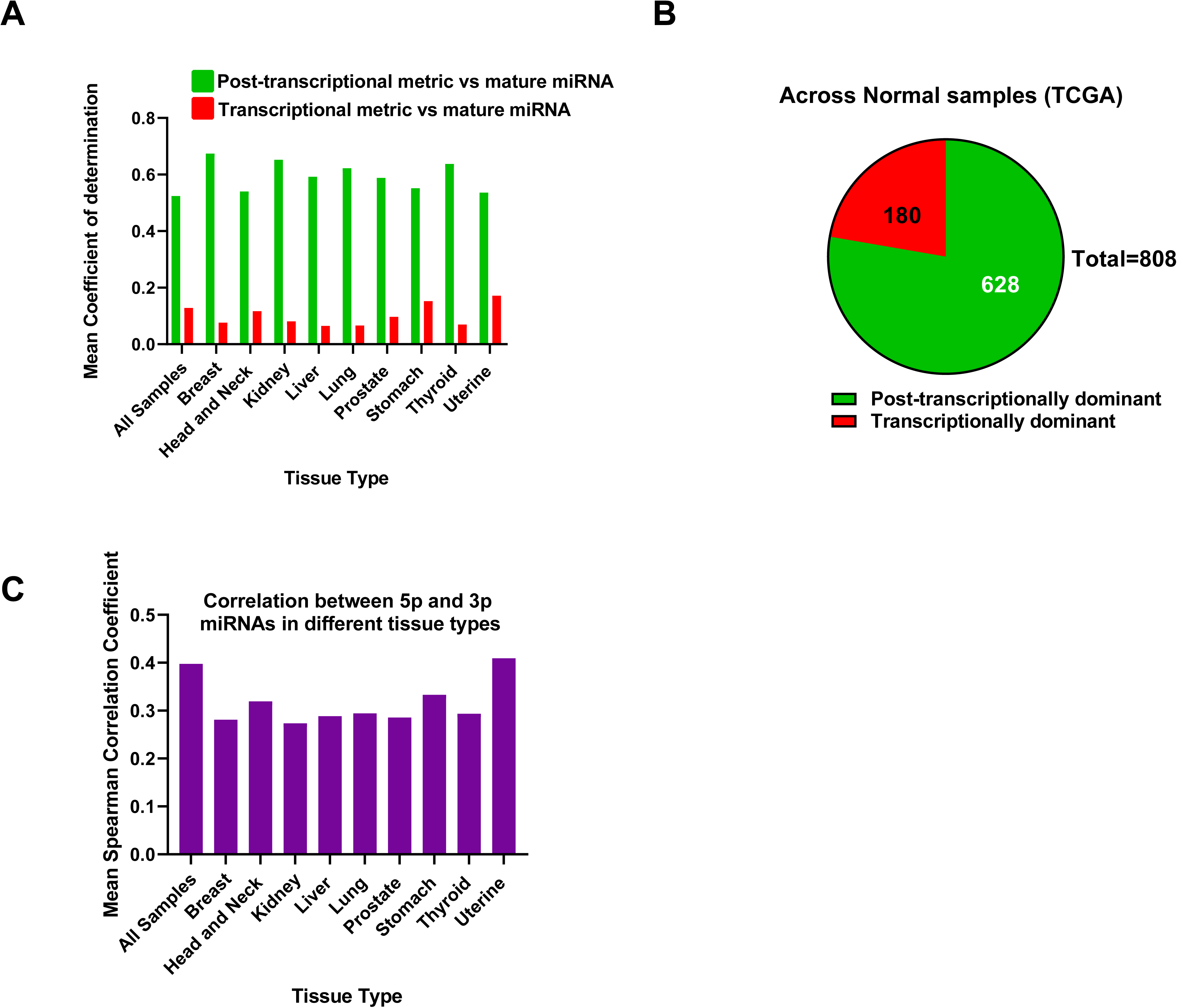
Post-transcriptional metrics is the major driver of the variation of mature miRNA expression in normal samples. **(A)** A bar graph plotting average contribution of post-transcriptional metrics to overall miRNA expression variation (green bars) and average contribution of transcriptional metrics (red bars) in each normal tissue type within TCGA. **(B)** A pie chart showing the number of transcription-dominant miRNAs and post-transcription-dominant miRNAs, calculated across all normal samples in the TCGA dataset. **(C)** A histogram showing mean Spearman correlation coefficients between 5p and 3p miRNAs (from the same hairpins) in each normal tissue type or across all normal samples in TCGA.

We next asked whether there are co-regulation patterns based on the post-transcriptional metric, the existence of which would suggest that groups of miRNAs undergo similar regulation at the post-transcriptional level. To address this, we calculated pair-wise correlation between miRNAs based on their post-transcriptional metrics, transcriptional metrics or mature miRNA expression levels. Indeed, similarity patterns emerge between miRNAs upon examination of the correlation matrices (Figure 6A). Some miRNAs, exemplified by those located in close genomic proximity with a clustered pattern, show stronger inter-miRNA correlation based on the transcriptional metric than post-transcriptional metric, which better explained correlation among mature miRNAs (see Group 2 in Figure 6A as an example). In contrast, other groups of miRNAs with better inter-miRNA correlation based on the post-transcriptional metrics could also be observed. One of such groups contained *let-7g* and miR-98, two let-7 family members known to be suppressed post-transcriptionally by LIN28B. Indeed, we observed that LIN28B is negatively associated with the post-transcriptional metrics of these miRNAs but not the transcriptional metrics (Figure 6B). Additionally, we noticed that there was a large co-regulated groups of miRNAs based on the post-transcriptional metric which were less correlated using the transcriptional metric (Group 1 in Figure 6A). To gain further insights into this group of post-transcriptionally co-regulated miRNAs, we calculated the average correlation between the post-transcriptional metrics of these miRNAs with mRNA genes, followed by gene set enrichment analyses. We observed that the post-transcriptional metrics of Group 1 miRNAs were significantly and positively associated with cell cycle, suggesting that these miRNAs could be co-regulated post-transcriptionally in proliferating cells (Figure 6C). In contrast, the transcriptional metrics of Group 1 miRNAs were not positively associated with cell cycle (Figure 6C). In addition to similarity across all cancer specimens in TCGA, the post-transcriptional metrics of miRNAs in Group 1 had stronger positive correlation among themselves in 28 out of 32 individual TCGA cancer types than the background level (Figure 6D), suggesting that these miRNAs are often co-regulated post-transcriptionally in many cancer types. The data above suggest that the co-clustering patterns of miRNAs on the post-transcriptional level could inform mechanisms of post-transcriptional regulation and related biological processes.

**Figure 6.**
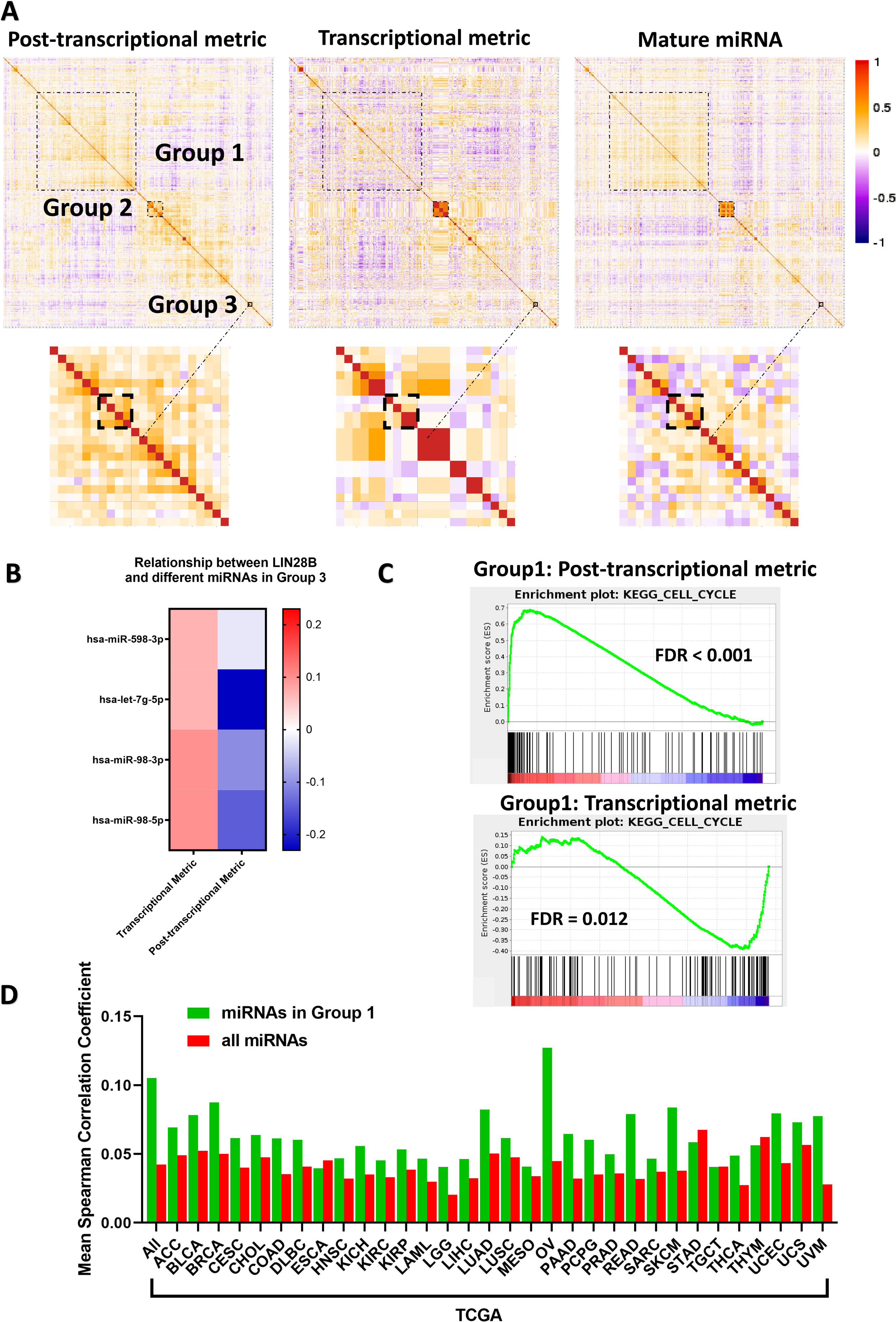
Co-regulation patterns based on post-transcriptional metrics reflect mechanisms of regulation and cancer processes. **(A)** Heatmaps are shown for correlation matrices based on post-transcriptional metrics, transcriptional metrics and mature miRNA expression. Spearman correlation was calculated based on all cancer samples in TCGA. Colors follow the color bar on the right. Specific groups of co-regulated miRNAs are indicated with dashed boxes. Enlarged sections of the heatmaps are shown on the bottom, focusing on group 3 miRNAs. **(B)** A heatmap shows Spearman correlation coefficients between LIN28B and post-transcriptional metrics or transcriptional metrics of miRNAs in Group 3. **(C)** Genes were ranked according to mean Spearman correlation coefficients with the post-transcriptional metrics (top) or transcriptional metrics (bottom) of miRNAs in Group 1. The resultant rank list was used to query the KEGG pathways in MSigDB database using GSEA. An enrichment plot is shown for the gene set KEGG_Cell_Cycle, which was the top KEGG gene set to be positively correlated with the post-transcriptional metrics of group 1 miRNAs. False discovery rates (FDR) are indicated. **(D)** Mean Spearman correlation coefficients were calculated among pairs of miRNAs within Group 1, using all TCGA cancer samples or within individual cancer types. Mean Spearman correlation coefficients between all miRNA pairs were also calculated as a background. Data were plotted with TCGA cancer types indicated.

## Discussion

How miRNAs are regulated and deregulated in disease is a long-standing question. Intuitively, one would think that the majority of the control occurs transcriptionally, because regulation at the transcriptional step is the most economical for the cell to avoid wasting building materials. In this study, we took advantage of the existing large public datasets in which both small and large RNAs have been profiled, and established transcriptional and post-transcriptional metrics to estimate the effects of transcriptional control and post-transcriptional regulation on mature miRNA levels. While these metrics may not be 100% perfect, because post-transcriptional regulation of the host genes themselves was difficult to be deduced from public data and thus challenging to be incorporated, two lines of evidence support that our estimates are close approximation of transcriptional and post-transcriptional regulation. First, we have examined several known regulatory relationships in which protein-coding genes regulate miRNA expression transcriptionally and post-transcriptionally, and our metrics reflect these known regulatory mechanisms to a good extent. Second, we have previously used the miRNA/host ratios (the post-transcriptional metric) to faithfully capture a number of rules of pri-miRNA processing, such as the optimal hairpin length and the CNNC motif promoting pri-miRNA processing, as well as the cooperation between optimal hairpin length and CNNC motif in pri-miRNA processing ^21^.

We found that the post-transcriptional metrics could explain the variance of mature miRNA expression more than twice that of the transcriptional metrics. Furthermore, the number of miRNAs that shows dominance by the post-transcriptional metric is also more than two times that of the transcription-dominant miRNAs. These data are consistent with the model that post-transcriptional regulation plays a more dominant role in shaping mature miRNA expression. Consistent with our findings, recent studies have shown that there are many proteins that can bind pre-miRNAs ^12, 47, 48^, some in a miRNA-specific fashion, thus potentially allowing the prevalence of post-transcriptional regulation to occur. Why more miRNAs would prefer post-transcriptional regulation? One possibility is that this mechanism creates diversity in miRNA expression pattern that is different from its host gene, thus utilizing the two gene products from the same transcript in potentially different ways. This possibility is suggested by our observation that post-transcription-dominance is particularly strong for broadly expressed host genes. Thus it may be more economical for cells to encode two different functions (host gene and miRNA) from the same transcript. Post-transcriptional regulation includes steps in DROSHA processing, DICER processing, AGO loading of miRNA duplex and stability of mature miRNAs. Which steps are the main contributors to post-transcriptional regulation in cancer and normal tissues? This question is difficult to answer at this moment, as existing large datasets do not have profiles of post-transcriptional regulatory intermediates, such as pre-miRNAs levels. Nevertheless, we observed that the ratios between 5p and 3p miRNAs from the same hairpin can be an important contributor. Some interesting miRNAs are affected, such as the tumor suppressor miR-146a, which is often deleted on human chromosome 5q in a subset of myelodysplastic syndromes ^45, 49^. The change in 5p/3p ratios may thus allow additional targeting from the often neglected star strand miRNA to play critical functional roles in some samples. Overall, we estimate that ∼20-30% of post-transcriptional regulation occurs at the step of 5p/3p ratio changes. It is possible that regulation at the DROSHA and DICER steps contribute to the remaining ∼70%. It is not fully clear what are the major processes that underlie miRNA 5p/3p ratio variation. It has been reported that miRNA target abundance may regulate miRNA stability ^50, 51^, which may contribute to this ratio variation. However, it is also possible that other steps in miRNA biogenesis, such as non-canonical processing, or currently unknown mechanisms, play important roles to regulate miRNA 5p/3p ratios in cancer and normal tissues.

Using the post-transcriptional metrics, we could group miRNAs by similarity, which could reflect co-regulation on the post-transcriptional level for miRNAs in the same group. We further demonstrate that the known regulatory relationship between LIN28B and let-7g and miR-98 could be read out through correlation of genes and the post-transcriptional metrics of co-regulated miRNAs. We postulate that similar strategies could be used to discover trans-regulatory mechanisms for miRNA processing, loading and stability.

## Method

### Source of Genomic Data

The table below shows the source of data and annotation, as well as the data download date in this study. Of note, we also performed analyses on data from earlier download dates, which showed similar results.

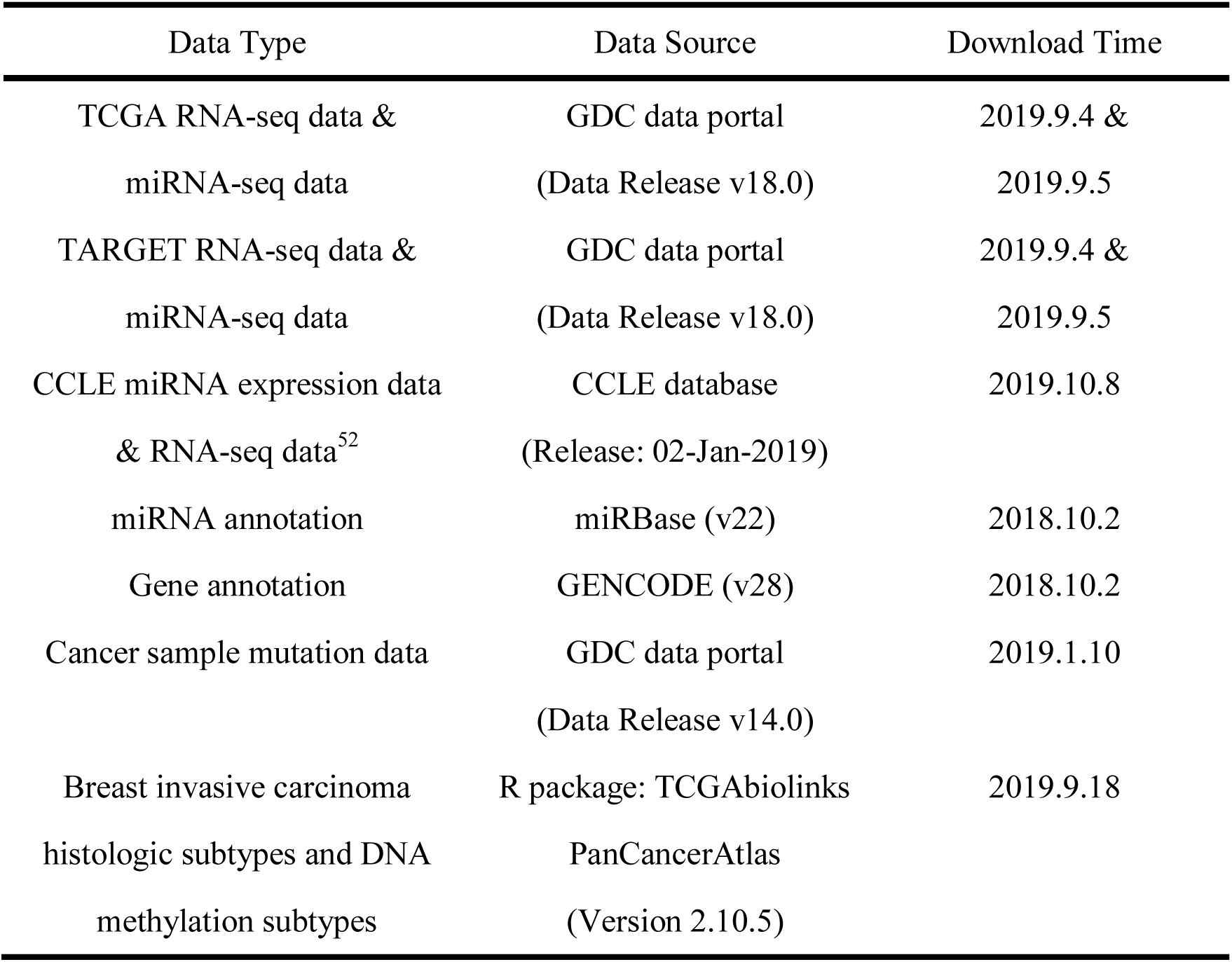

### Pairing of miRNAs and Host Genes

For each mature miRNA in miRBase, we identified its host gene using the miRBase-annotated genomic location. Host genes were identified using the gene annotation from GENCODE ^53^, satisfying the condition that a given miRNA is located completely within the gene body (including introns) of the GENCODE gene. To avoid ambiguity, miRNAs which could be mapped to two or more host genes, or could be produced from two or more genomic locations were removed from analysis. Thus, we have one-to-one miRNA/host gene pairs for all remaining miRNAs.

### Data Normalization and Filtering

Data for miRNA and mRNA expression were normalized by custom scripts with a method similar to that in the DESeq2 R package) ^54^. The calculation consists two steps:

1. For each sample, we calculated an adjustment factor. Only genes which are expressed in all samples (absolute read counts are more than zero in every sample) were considered in the calculation of adjustment factors. The calculation equation is:

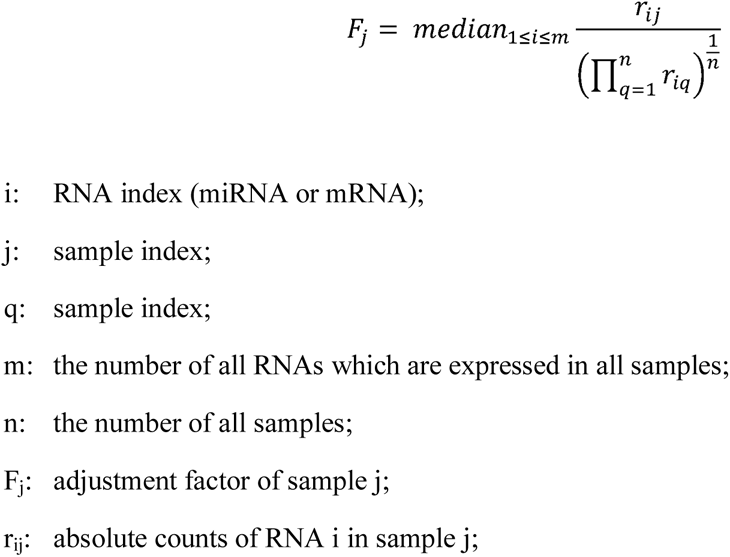

2. For each RNA (miRNA or mRNA) in each sample, we used the adjustment factor to adjust the count to obtain the normalized count.

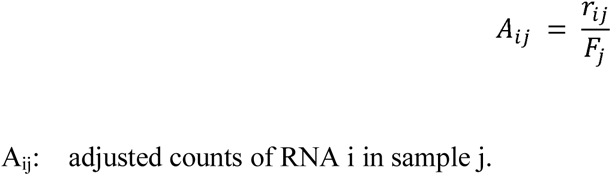

For miRNA, we used these normalized counts as the expression level. For mRNA, the normalized count was further divided by the length of the corresponding mRNA, which was then used as the expression level.

In addition to the above normalization, we also used normalized data obtained from GDC data portal, with miRNA expression quantified in RPM, and mRNA expression quantified in FPKM. Similar results were obtained regardless of which normalization method was used.

Filtering of miRNA/host gene pairs: we filtered out those pairs whose miRNA expression or host gene expression was consistently low across almost all samples in any given dataset. Specifically, we used the following criteria to determine which miRNA/host gene pair to retain for analysis: (1) the absolute read count of the miRNA is greater or equal to 8 in at least 20 samples; and (2) the absolute read count of the host gene is greater or equal to 8 in at least 20 samples. Unless specified otherwise, all retained miRNA/host gene pairs were used for analysis.

### Development of Transcriptional and Post-Transcriptional Metrics

For each miRNA, we used its miRNA/host-gene ratio as its post-transcriptional metric and the expression level of its host gene as its transcriptional metric.

When calculating the miR/host ratio, we added a small value, to avoid mathematical artifacts from zero numbers, based on the following formula:

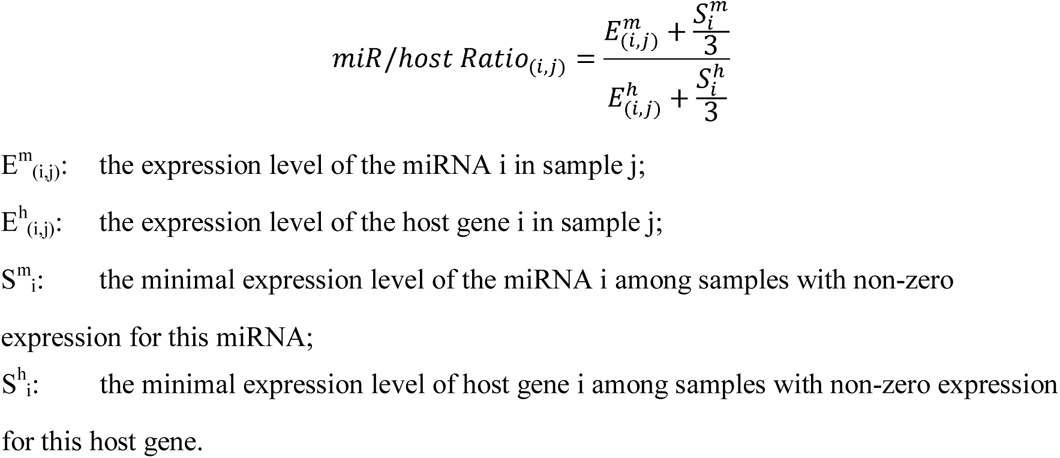

Three known cases were used to verify our transcriptional metrics and post-transcriptional metrics. Spearman correlation between the transcriptional metric (or post-transcriptional metric) of hsa-let-7g-5p and the expression level of LIN28B was calculated, using all cancer samples in TCGA, and the statistically significance of the Spearman correlation was calculated (using R function cor.test). For MYC regulating hsa-miR-17-5p, hsa-miR-18a-5p, hsa-miR-19a-3p and hsa-miR-20a-5p, we performed similar Spearman correlation calculations between the expression levels of MYC and the metrics of those miRNAs. For p53 regulating hsa-miR-34a-5p, we compared the transcriptional metrics (or post-transcriptional metrics) of samples that were annotated as wildtype for p53 or mutant for p53 (according to GDC data portal), and performed one-tailed Wilcoxon rank sum and signed rank tests (using R function cor.test).

### Estimating Contribution of Transcriptional and Post-Transcriptional Regulation to Mature miRNA Expression

Coefficient of determination (R^2^) was calculated to assess the contribution of transcriptional regulation and post-transcriptional regulation on the expression variation of miRNAs. We obtained coefficients of determination by computing the Pearson correlation coefficients between mature miRNA expression and the corresponding transcriptional metric (or post-transcriptional metric) and squaring the result R. Calculation was performed based on all cancer samples (or normal samples) in TCGA, or specifically within samples of a given cancer type. Mean coefficient of determination of all miRNAs was used to represent the overall influence of post-transcriptional metric or transcriptional metric on mature miRNA expression. For analyzing individual cancer types in TCGA, we removed miRNA/host gene pairs whose absolute reads counts are non-zero in fewer than 10 samples in the corresponding subset.

Individual miRNAs were categorized as transcription-dominant or post-transcription-dominant by comparing Pearson correlation between mature miRNA expression and transcriptional or post-transcriptional metrics. Dominance was assigned to the larger correlation coefficient.

For samples from normal tissues of TCGA database, TARGET database, and CCLE database, we performed similar analyses. For CCLE dataset analyses, we removed miRNAs whose microarray expression levels are background values in > 20% of samples and genes whose adjusted count values are background values in > 20% of samples.

### Analysis of miRNAs with Broadly Expressed Host Genes

We divided miRNAs into two groups by whether their host genes are broadly expressed. Broadly expressed host genes were defined as those with absolute read counts greater than or equal to 8 in every sample within all TCGA and TARGET samples. Then, we compared the number of post-transcriptionally dominant miRNAs to transcriptionally dominant miRNAs between these two groups. P-value was calculated by Fisher exact test.

### Cancer Type and Subtype Classification

We used the same number of variables for host gene, miRNA/host ratio and mature miRNAs for prediction. Because some miRNA hairpins produce both 5p and 3p miRNAs, we thus only took the more abundantly expressed miRNA from each hairpin for analysis. For mature miRNAs located in genomic clusters, we again only selected the most abundantly expressed miRNA from the corresponding cluster for use in prediction analyses. In total, 493 miRNA/host gene pairs were retained after the above processing.

In addition to using all 493 miRNA/host gene pairs in the perditions (**Supplementary Figure S5**), we also performed filtering to focus on informative miRNA/host gene pairs. The filtering was performed by selecting both housekeeping miRNAs and tissue-enriched miRNAs. We defined housekeeping miRNAs as miRNAs which are expressed (i.e. having non-zero read count) in over 98 percent of all samples. For any cancer type or subtype, we defined tissue-enriched miRNAs based on two criteria: (1) miRNAs having non-zero expression level in > 98% of samples in this cancer type; and (2) miRNAs showing higher expression in 75% of samples in this cancer type than the median expression in all TCGA cancer samples. Host genes were also similarly defined as housekeeping and tissue-enriched. Only miRNA/host gene pairs that satisfy filtering on both miRNA and host genes were selected for prediction analyses (a total of 117 pairs for cancer type prediction; a total of 96 pairs for breast cancer histology subtype prediction; a total of 96 pairs for breast cancer methylation subtype prediction).

For each type of metric (transcriptional, post-transcriptional and mature miRNAs), we applied supported vector machine (SVM) to classify cancer type/subtype. R package e1071 and caret was used in building and parameter tuning of SVM ^55, 56^. We used 10-fold cross-validation to compare the prediction performance (using Macro-F1 scores).

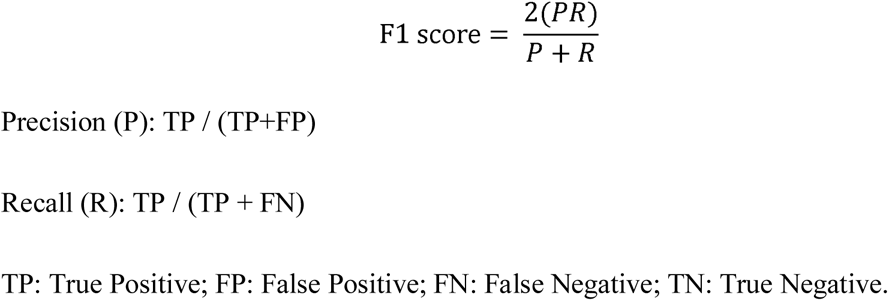

The Macro-F1 score was calculated as the average of F1 scores, with each F1 score calculated for each cancer type (or subtype).

We used 10-fold cross-validation to compare prediction accuracy. To perform each 10-fold cross-validation, we randomly divided all samples into 10 equal fractions, with each fraction containing 10% of samples from each cancer type. For each predication, we used one of these data fractions as the test set and all remaining data fractions as the training set. We trained SVM using the training set and calculated the Macro-F1 scores of the model on the test set. In total, 10 predictions were performed, with each one based on one 10% fraction of the data for testing and the rest for training. For cancer type prediction in TCGA, SVM-based multi-class predictor models were used, so that samples were predicted to be one of the 32 types of cancers. For breast cancer histology subtype prediction, we used samples defined as breast invasive ductal carcinoma, breast invasive lobular carcinoma, breast mixed ductal and lobular carcinoma and an “other” category (containing other breast cancer subtypes). We then built a 4-class predictor to classify each sample into one of these three subgroups. The difference compared to cancer type prediction is that we performed 10-fold cross-validation four times and combined the results. For breast cancer methylation subtype prediction, there were four known subgroups c1, c2, c3 and c4 ^34^. We thus built a 4-class predictor to classify each sample into one of these four groups. We performed 10-fold cross-validation twice and combined the results. For those analysis without secondary filtering, we performed 10-fold cross-validation twice for prediction on cancer type, ten times for breast cancer histology subtype prediction and twice for breast cancer methylation subtype prediction.

For predictions based on 5p/3p miRNA ratios, we used a similar approach with 10-fold cross-validations twice for cancer type prediction based on SVM. For random prediction controls, we randomly scrambled the cancer type assignment.

Statistical significance between different metrics was calculated by one-tailed Wilcoxon matched-pairs signed rank test.

### Data Clustering

We used principle component analysis to extract principle components from post-transcriptional metrics in TCGA and TARGET datasets. We then clustered the samples using hierarchical clustering based on the Top 100 principle components. Hierarchical clustering was performed using Euclidian distance and complete linkage. To avoid overcrowding of the dendrogram, we also performed hierarchical clustering on a subset of samples. These samples were selected based on the following: fifty samples randomly selected from TCGA-LAML dataset, fifty samples randomly selected from TARGET-AML dataset, and 200 non-AML samples randomly selected from TCGA or TARGET.

### Analysis of miRNA 5p-3p Ratio

For analysis on 5p-3p ratio, we only used 5p-3p pairs whose 5p isoforms and 3p isoforms both have >= 8 absolute read counts in at least 20 samples. For analysis in individual cancer types, we filtered 5p-3p pairs in a similar way by determining which 5p-3p pairs to retain using the following criterion: 5p miRNA, 3p miRNA and the corresponding host gene all have non-zero absolute read counts in at least 10 samples of this cancer type.

For each 5p/3p pair in one sample, we calculated 5p/3p ratio as below:

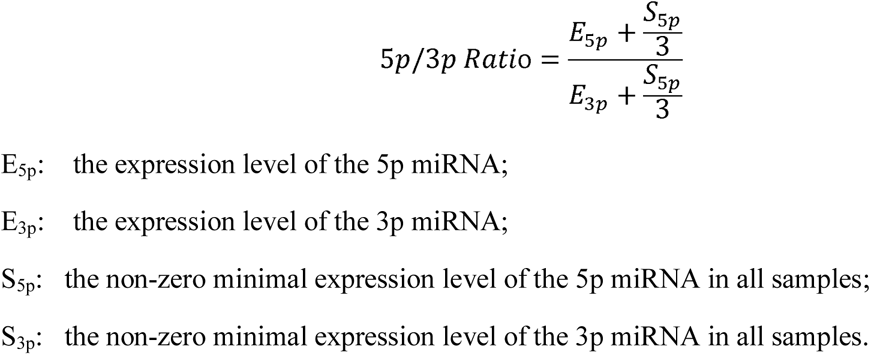

Histogram of 5p/3p ratios of hsa-mir-146a was drawn in R; a smooth density estimate was also added. We finished this figure with ggplot2 and R ^57^.

We also used 5p/3p ratios to explain the post-transcriptional regulation of miRNAs. Then, we calculated the Pearson correlation coefficients and use R^2^ between 5p/3p ratios and miR/host ratios in all TCGA cancer samples, or within individual cancer types.

### Analysis of Co-regulated miRNAs

We calculated Spearman correlation coefficients between every pair of miRNAs based on their post-transcriptional metrics, transcriptional metrics or mature miRNA expression levels. Correlation matrix was clustered with hierarchical clustering using 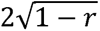 as the distance measure (r: Spearman correlation coefficient calculated between post-transcriptional metrics of two miRNAs) and average linkage. Three heatmaps were plotted based on correlation matrices of post-transcriptional, transcriptional and mature miRNA expression. To assist direct comparison, the order of miRNAs in these heatmaps were based on the sample order of the hierarchical clustering which were based on clustering of the post-transcriptional metrics.

To identify which genes or pathways are associated with a group of co-regulated miRNAs, for each mRNA gene, we calculated the mean Spearman correlation coefficient between this gene and every miRNA within the group. Genes were then sorted based on mean correlation levels. GSEA analysis (pre-ranked mode) was then performed to identify significantly enriched gene sets (GSEA 3.0).

### Statistical Methods

We performed statistical analysis, data collection, data processing, and figure plotting using custom codes in R (R version 3.4.1) and Perl (Perl 5, version 16), as well as using GraphPad Prism. For calculation of Spearman correlation coefficient, we did log-transformation before the calculation. If there were zero values in a data series before log-transformation, we add one third of the non-zero minimal value of this data series.

## Author Contribution

J.L. conceived and supervised the study. D.Z. performed all bioinformatics analysis. C.R. initiated the use of the post-transcriptional metric. J.T. assisted with data interpretation. D.Z. and J.L. wrote the manuscript.

## Acknowledgment

The results published here are in part based upon data generated by the TCGA Research Network: https://www.cancer.gov/tcga. The TCGA data used for this analysis are available at https://portal.gdc.cancer.gov/projects. The results published here are in part based upon data generated by the Therapeutically Applicable Research to Generate Effective Treatments (https://ocg.cancer.gov/programs/target). The TARGET data used for this analysis are available at https://portal.gdc.cancer.gov/projects. Research in this study is supported in part by NIH grant R01GM116855 (J.L.).

