## Supplemental Figures for "Post-transcriptional Regulation is the Major Driver of microRNA Expression Variation"

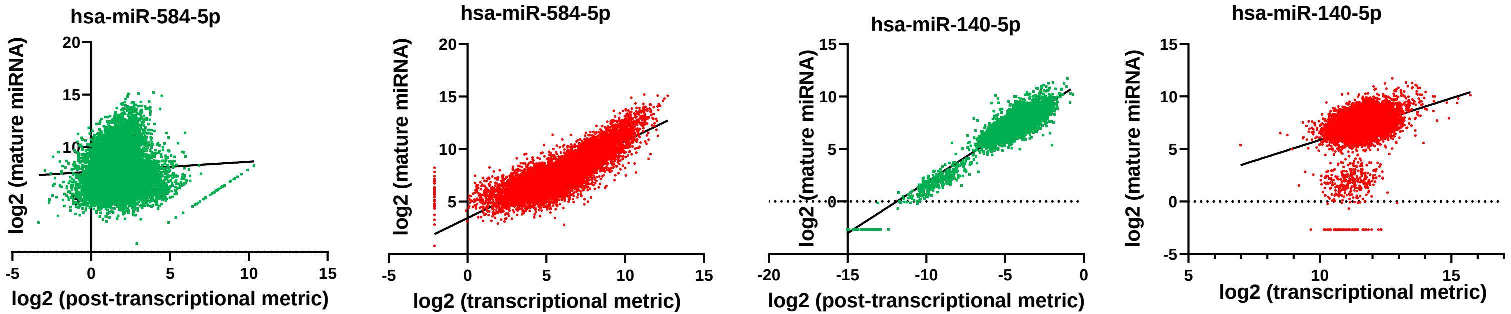

hsa-miR-584-5p
log2 (mature miRNA)
log2 (post-transcriptional metric)
hsa-miR-584-5p
log2 (mature miRNA)
log2 (transcriptional metric)
hsa-miR-140-5p
log2 (mature miRNA)
log2 (post-transcriptional metric)
hsa-miR-140-5p
log2 (mature miRNA)
log2 (transcriptional metric)

**Figure S1. Examples of miRNAs whose expression correlate with the transcriptional or post-transcriptional metrics.** For hsa-miR-584-5p and hsa-miR-140-5p, scatter plots show the relationship between mature miRNA expression and transcriptional (red) or post-transcriptional (green) metrics across all cancer samples in TCGA. Each dot reflects one sample. Fitted lines are shown based on linear regression.

A

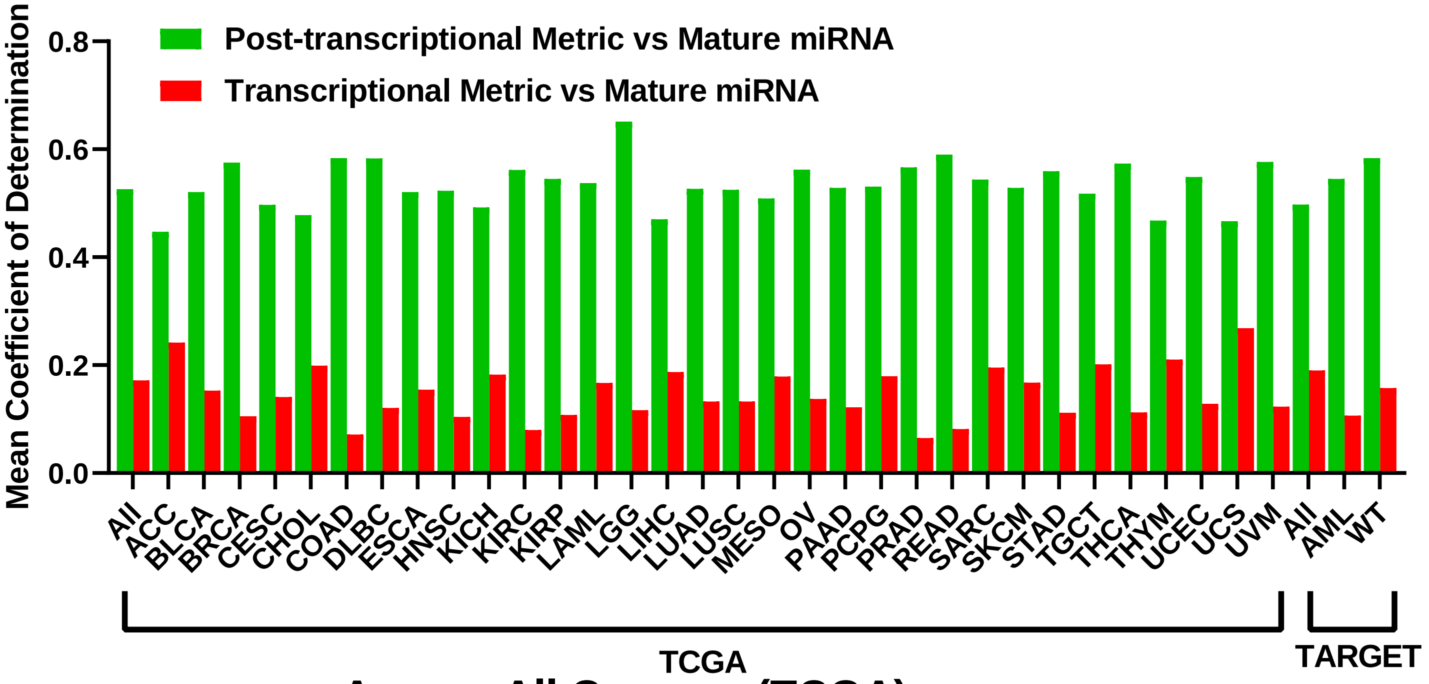

Post-transcriptional Metric vs Mature miRNA
Transcriptional Metric vs Mature miRNA
Mean Coefficient of Determination
0.8
0.6
0.4
0.2
0.0
All ACC BLCA BRCA CESC CHOL COAD DLBC ESCA HNSC KICH KIRC KIRP LAML LGG LIHC LUAD LUSC MESO OV PAAD PCPG PRAD READ SARC SKCM STAD TGCT THCA THYM UCEC UCS UVM All AML WT
TCGA
TARGET


B

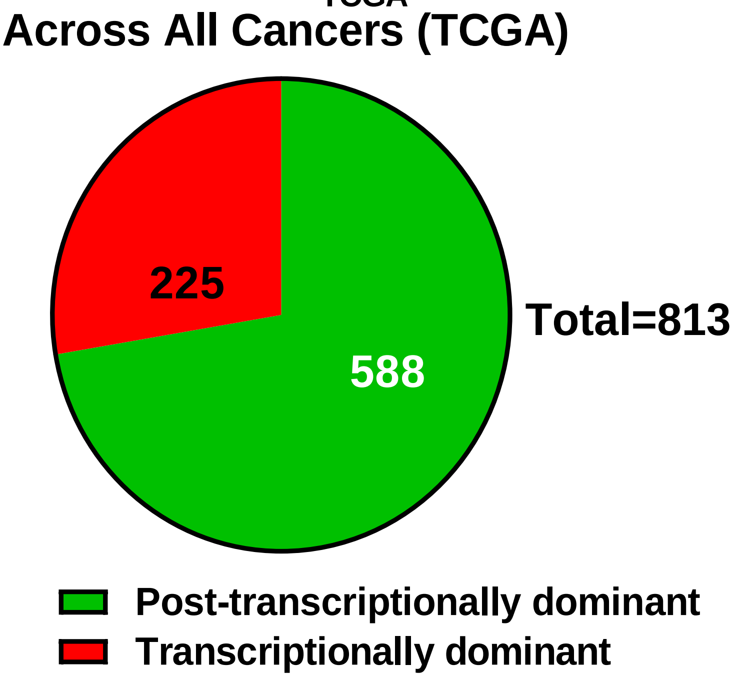

Across All Cancers (TCGA)
225
588
Total=813
Post-transcriptionally dominant
Transcriptionally dominant


C

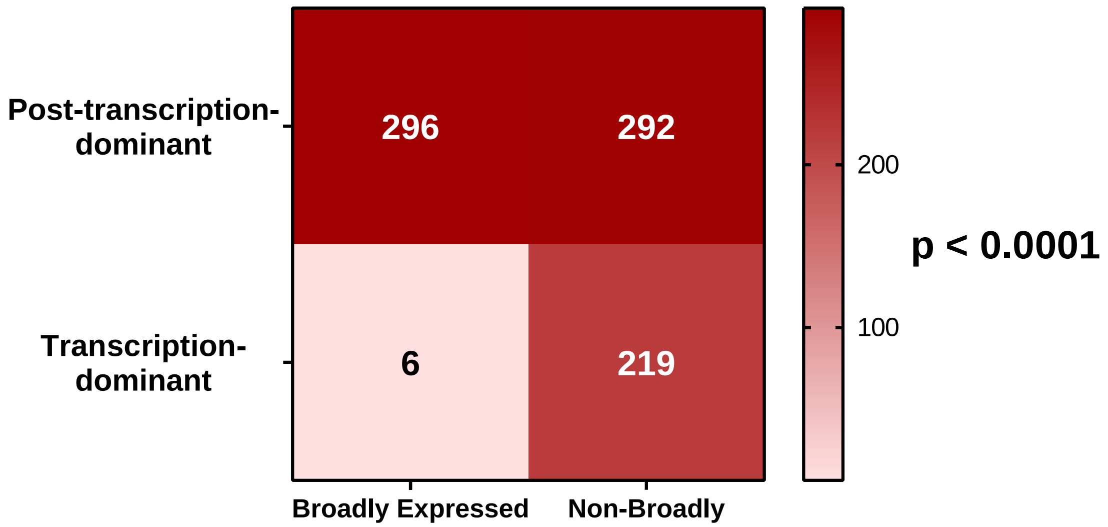

Post-transcription-dominant
Transcription-dominant
296
292
6
219
200
100
p < 0.0001
Broadly Expressed
Non-Broadly

**Figure S2. Post-transcription-dominance reflected in an alternative method of data normalization.** We used normalized data downloaded from GDC data portal in our calculation. In such data, FPKM values were used to reflect mRNA gene expression level and RPM values were used to reflect mature miRNA expression levels. **(A)** A bar graph plotting average contribution of post-transcriptional metrics to overall miRNA expression variation (green bars) and average contribution of transcriptional metrics (red bars) in each cancer type or across all cancer samples is shown. Sample sources (TCGA and TARGET dataset) are indicated. **(B)** A pie chart showing the number of transcription-dominant miRNAs and post-transcription-dominant miRNAs, calculated across all cancer samples in the TCGA dataset. **(C)** miRNAs were separated into those with broadly-expressed host gene patterns and those with non-broadly-expressed host gene patterns. These two groups were further separated based on transcription-dominance or post-transcription-dominance. Numbers indicate the number of miRNAs, with colors reflecting the numbers. A color bar is shown on the right. P value based on Fisher exact test is indicated.

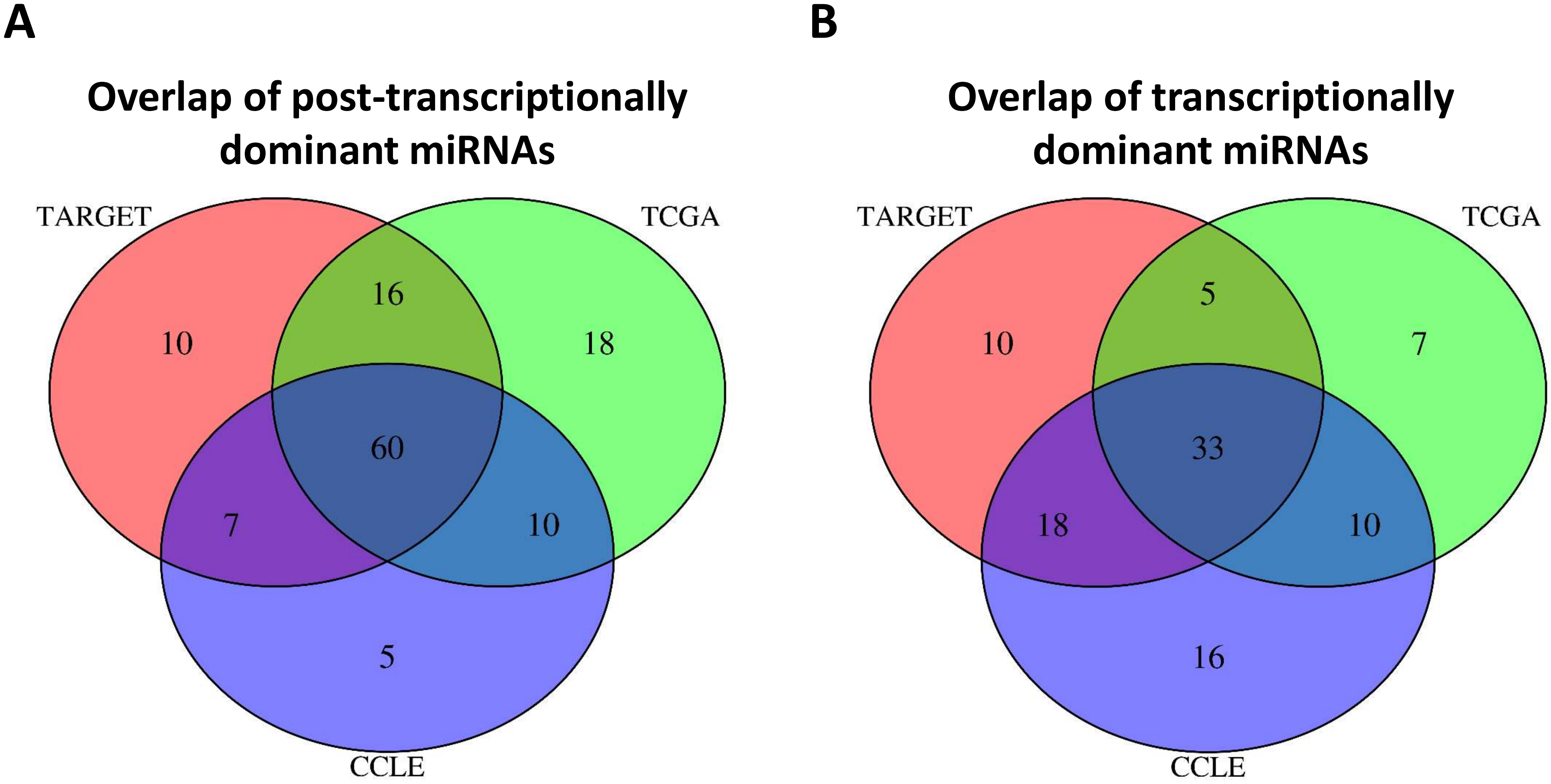
A
Overlap of post-transcriptionally dominant miRNAs
TARGET
TCGA
CCLE
10
16
18
60
7
10
5
B
Overlap of transcriptionally dominant miRNAs
TARGET
TCGA
CCLE
10
5
7
33
18
10
16

**Figure S3. Transcription-dominant and post-transcription-dominant miRNAs are similar across datasets. (A)** A Venn diagram showing the overlap among post-transcription-dominant miRNAs identified separately based on all cancer samples in TARGET dataset, TCGA dataset, and CCLE dataset. Numbers indicate the number of miRNAs. Only miRNAs and host genes that passed filtering in all three datasets were analyzed. **(B)** A Venn diagram showing the overlap among transcription-dominant miRNAs identified separately based on all cancer samples in TARGET dataset, TCGA dataset, and CCLE dataset.

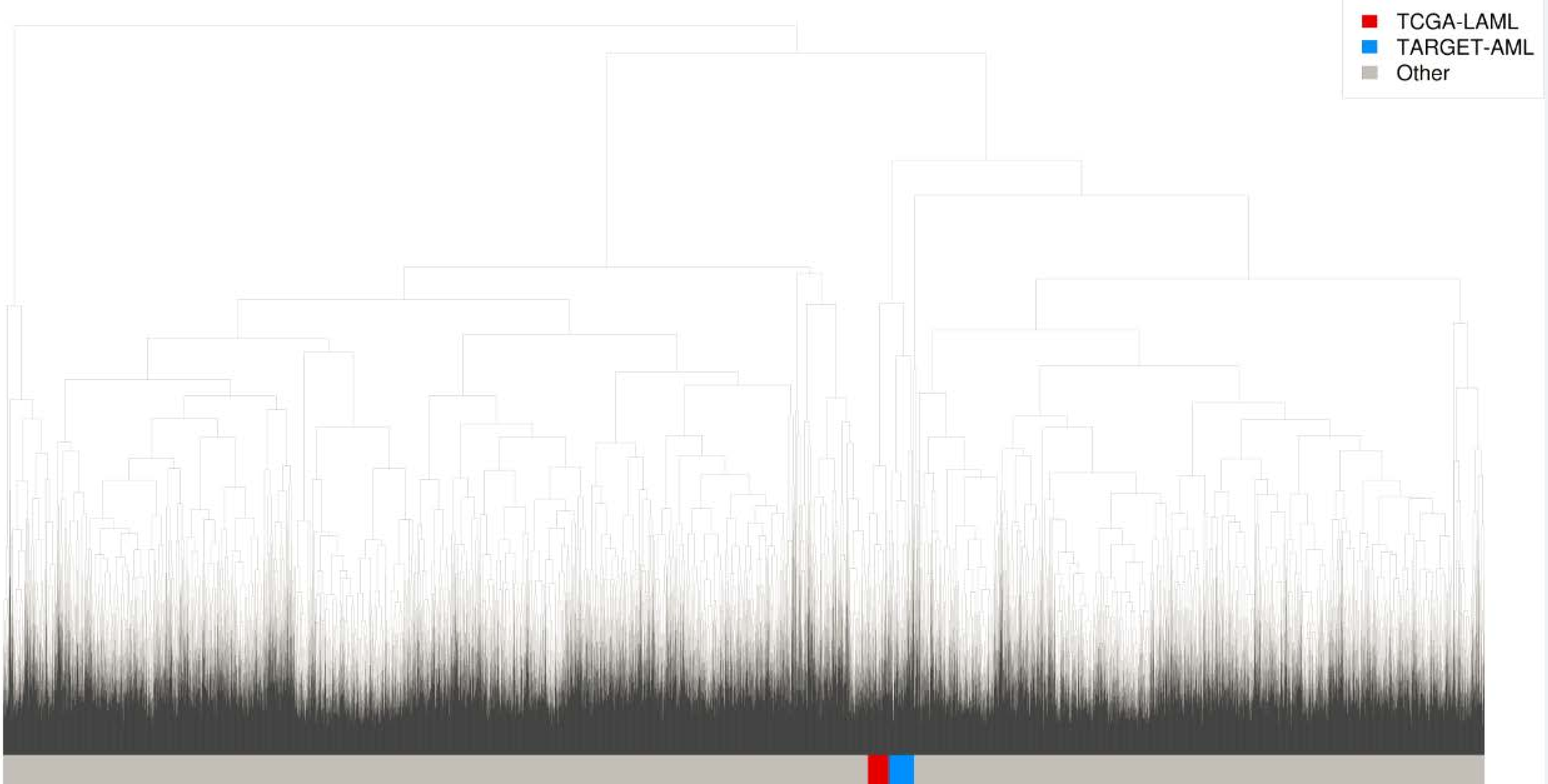
TCGA-LAML
TARGET-AML
Other

**Figure S4.** Hierarchical clustering was performed based on the post-transcriptional metrics of miRNAs on all cancer samples in TCGA and TARGET AML samples. The dendrogram from the clustering is shown with AML samples from TCGA and TARGET datasets indicated.

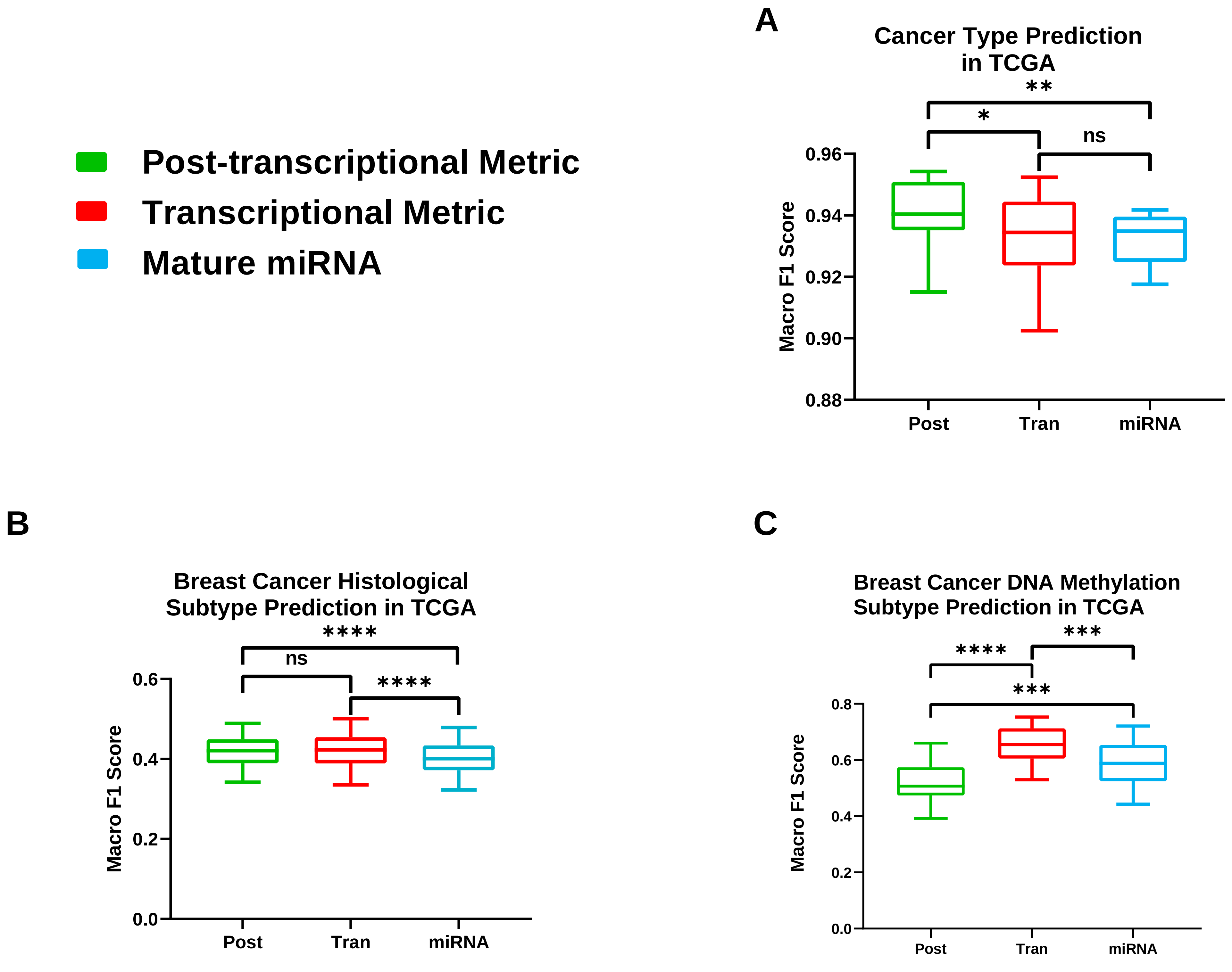
Post-transcriptional Metric
Transcriptional Metric
Mature miRNA
A
Cancer Type Prediction in TCGA
Macro F1 Score
0.96
0.94
0.92
0.90
0.88
Post
Tran
miRNA
**
*
ns
B
Breast Cancer Histological Subtype Prediction in TCGA
Macro F1 Score
0.6
0.4
0.2
0.0
Post
Tran
miRNA
****
ns
****
C
Breast Cancer DNA Methylation Subtype Prediction in TCGA
Macro F1 Score
0.8
0.6
0.4
0.2
0.0
Post
Tran
miRNA
****
***
***

**Figure S5. Additional data on the superiority of post-transcriptional metrics in prediction.** 493 miRNA/host gene pairs were used in prediction (see Methods). **(A)** SVM-based multi-class predictors were developed based on the indicated metrics to predict cancer types in TCGA. Each test sample was classified into one of the 32 cancer types. Predication accuracies are plotted using box-and-whisker plot, with data representing results from multiple 10-fold cross-validation. **(B)** SVM-based multi-class predictors were developed to predict breast cancer histologic subtypes. Prediction accuracies are shown. **(C)** SVM-based multi-class predictors were developed to predict breast cancer methylation subtypes. Prediction accuracies are shown. (ns: not significant; * $p < 0.05$; ** $p < 0.01$; *** $p < 0.001$; **** $p < 0.0001$).
